# An evolutionarily conserved mechanism for production of secondary meristems in land plants

**DOI:** 10.1101/555607

**Authors:** Yukiko Yasui, Shigeyuki Tsukamoto, Tomomi Sugaya, Ryuichi Nishihama, Quan Wang, Katsuyuki T. Yamato, Hidehiro Fukaki, Tetsuro Mimura, Hiroyoshi Kubo, Klaus Theres, Takayuki Kohchi, Kimitsune Ishizaki

## Abstract

A variety of plants in diverse taxa can reproduce asexually via vegetative propagation, in which clonal propagules with new meristem(s) are generated directly from vegetative organs. A basal land plant, *Marchantia polymorpha*, develops clonal propagules, gemmae, in a specialized receptacle, gemma cup. Here we report an R2R3-MYB transcription factor, designated GEMMA CUP-ASSOCIATED MYB 1 (GCAM1), which is an essential regulator of gemma cup development in *M. polymorpha*. Although gemma cups are a characteristic gametophyte organ for vegetative reproduction in a taxonomically restricted group of liverwort species, phylogenetic and interspecific complementation analyses supported the orthologous relationship of GCAM1 to regulatory factors for axillary meristem formation, e.g. Arabidopsis RAXs and tomato Blind, in angiosperm sporophytes. The present findings in *M. polymorpha* suggest an ancient acquisition of a regulatory mechanism for production of secondary meristems, and the use of the mechanism for diverse developmental programs during land plant evolution.

## Introduction

The plastic nature of plant architecture can be traced back to the activity of meristems, which are pools of pluripotent cells located at the tips of growing plant bodies. In seed plants, the primary shoot apical meristem (SAM) is established during embryonic development. Following germination, secondary meristems are continuously produced in the axils of leaves, and form the basis of branching in flowering plants. Many plant species also develop ectopic meristems at different positions of the plant body^1, 2^.

Vegetative reproduction, a form of asexual reproduction in plants, is a developmental process by which clonal progeny arise directly from parental tissues. This process is based on the remarkable potential of plants to proliferate meristems, which develop whole plantlets from differentiated tissues^1, 3, 4^. Diverse plants have the capability for vegetative reproduction, which occurs naturally from a variety of vegetative tissues, e.g. stems (rhizomes: asparagus, iris, ginger, bulbs: onion, tubers: potato, corms: crocus, and runners: strawberry, wild mint), roots (sweet potato), and leaves (*Kalanchoe*), or can be induced artificially by means of various techniques, including cuttings, grafting, and tissue culture. Vegetative reproduction is considered to be an important strategy for efficient plant propagation and survival in the natural environment^5^, as well as for agriculture and horticulture^6^.

The liverwort, *Marchantia polymorpha* L., as well as certain related species in the class Marchantiopsida, develops specialized organs for vegetative reproduction, termed the gemma cup or cupule^7^. Clonal propagules, called gemmae, develop from single epidermal cells at the base, or floor, of gemma cups. Thus, the floor cells have the capability to produce clonal plantlets. Gemma cups are formed periodically along the dorsal midrib of a thallus. The restricted occurrence of gemma cups along the midrib suggests that they originate from the dorsal merophyte above the apical cell^7^. Over 100 gemmae can develop in a single gemma cup. The development of gemma cups has been well described on the basis of a series of histological observations on *M. polymorpha*^8^. According to the detailed observations by Barnes and Land^8^, the precursors of the floor epidermis in a gemma cup are recognizable as close as the third cell back from the apical cell on the dorsal surface. The floor epidermis is a region where periclinal cell divisions to generate protodermal and subprotodermal cell layers of the air chamber are suppressed, and consecutive elongate epidermal cells are instead observed. The single-layered epidermal cells comprise the gemminiferous region, an area that will become the basal epidermis of a gemma cup. These cells undergo repeated anticlinal divisions to enlarge the area of the gemma cup floor, allowing for increase proliferation of gemma production. Some gemma cup floor cells begin to develop gemmae from an early stage of gemma cup development, and growth of the gemma cup occurs concomitantly with gemma development and maturation (Supplementary Figure 1). The frequency of gemma cup formation is influenced by a variety of environmental factors, such as light and nutrients^9,10,11^. However, the molecular mechanisms underlying gemma and gemma cup development are largely unknown.

*Marchantia polymorpha* is a member of an early diverging lineage among land plants. The species offers a number of advantages for genetic studies: low genetic redundancy^12^, a haploid-dominant life cycle, and practicable genetic tools^13^, such as high-efficiency transformation^14,15^ and gene modification techniques^16,17,18,19^. Recent molecular genetic studies on bryophytes, including *M. polymorpha*, have revealed several key developmental regulators for the gametophyte generation that have orthologous counterparts in angiosperms shown to control analogous, but not homologous, developmental processes in the sporophyte generation^20,21,22,23,24,25^. These observations suggest that a considerable number of developmental regulatory modules were acquired in the common ancestor of land plants before the divergence of bryophytes and vascular plants^26,27^.

In the present study, we undertook a comprehensive transcriptome analysis and isolated a R2R3-MYB transcription factor gene, *GCAM1*, which was predominantly upregulated in developing *M. polymorpha* gemma cups. Molecular and genetic characterization revealed a critical role for GCAM1 in gemma cup development. Cell proliferation without tissue differentiation was observed in plants ectopically expressing *GCAM1*. Although the gemma cup is a characteristic gametophytic organ for vegetative reproduction in certain species of Marchantiopsida, phylogenetic and interspecific complementation analyses supported the orthologous relationship of *GCAM1* with angiosperm genes of the R2R3-MYB subfamily 14^28^, which are involved in axillary meristem formation. Potential mechanisms shared between vegetative reproduction in the gametophyte of liverworts and axillary meristem formation in the sporophyte of angiosperms are discussed.

## Results

### An R2R3-MYB transcription factor is upregulated in the gemma cup

To screen for potential key regulators involved in vegetative reproduction in *M. polymorpha*, we performed RNA-seq analyses comparing the transcriptome of gemma cups with that of the entire young thallus yet to form gemma cups and that of the midrib region (Fig. 1a). We obtained more than 14 × 10^6^ reads from each of the samples (triplicates per tissue; Supplementary Table 1). More than 90% of the reads per library were mapped to the *M. polymorpha* genome sequence version 3.1^12^. Genes for which the reads per kilobase per million (RPKM) value was > 1 were considered to be ‘expressed’. Among 19,287 annotated genes, 11,699 genes (61%) were expressed in the gemma cup. We identified 1391 and 1483 genes with greater than 2-fold changes in the gemma cup compared with their expression in the young thallus and midrib tissue, respectively. A total of 664 genes were upregulated in both comparisons (Fig. 1b and Supplemental Data Sets 1–3). Of the 664 genes, 10 were annotated as transcription factors (Supplementary Table 2).

**Fig. 1.**
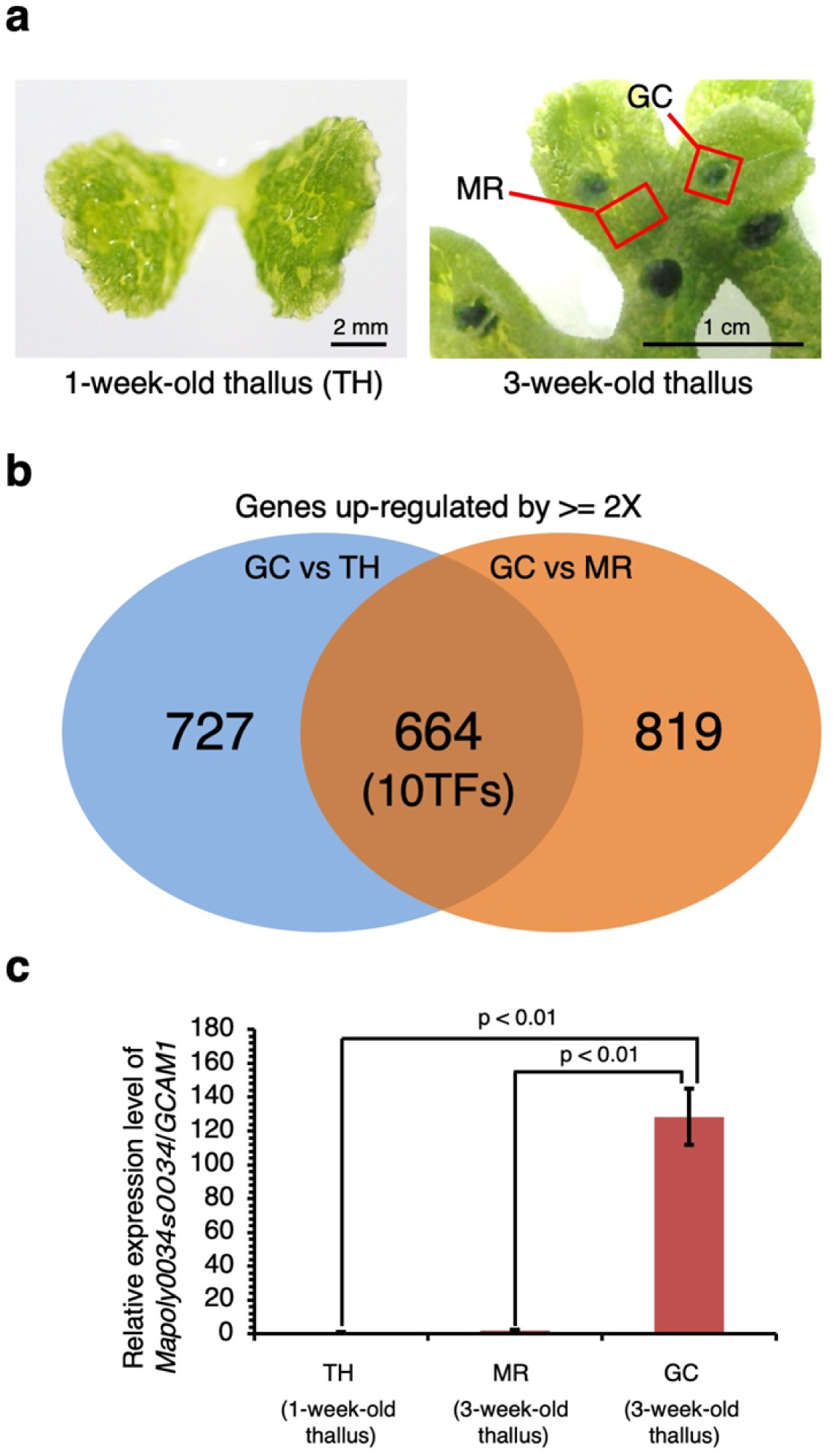
Identification of genes preferentially expressed in the gemma cup of *Marchantia polymorpha*. **(a)** Tissues used for mRNA-seq analysis. GC, gemma cup; TH, 1-week-old thallus; MR, midrib. **(b)** Venn diagram of genes upregulated by more than two-fold in the gemma cup. **(c)** qRT-PCR analysis of Mapoly0034s0034/*GCAM1* in vegetative tissues. The elongation factor 1a (Mp*EF1a*) gene was used as a reference (Welch’s *t*-test, *p* < 0.01, *n* = 3).

Among the transcription factors, we focused on Mapoly0034s0034, a gene encoding an R2R3-MYB transcription factor. The RPKM value of this gene in the gemma cup was 82.1, which represented 100-fold and 13-fold upregulation compared with expression in the young thallus and the midrib, respectively (Supplementary Table 2). Quantitative reverse-transcription PCR (qRT-PCR) analysis confirmed that Mapoly0034s0034 transcripts accumulated at a significantly higher level in gemma cups than in the young thallus and midrib tissue (Fig. 1c). We designated the *Mapoly0034s0034* gene as *GEMMA CUP-ASSOCIATED MYB 1* (*GCAM1*). Genetic nomenclature is as outlined in Bowman *et al*.^29^.

Next, we generated transgenic *M. polymorpha* plants expressing a β-glucuronidase (GUS) reporter gene under the control of the *GCAM1* promoter (*GCAM1pro:GUS*). We generated a construct in which the GUS reporter gene was translationally fused to a *GCAM1* genomic fragment, containing 5.2-kb upstream of the ATG start codon. In *GCAM1pro:GUS* transgenic lines, histological GUS staining was detected in the apical notches of the 10-day-old thallus (Fig. 2a). In mature thalli with gemma cups, significant promoter activity was detected in the apical notches, the floor of gemma-cups, and developing gemmae (Fig. 2b–d).

**Fig. 2.**
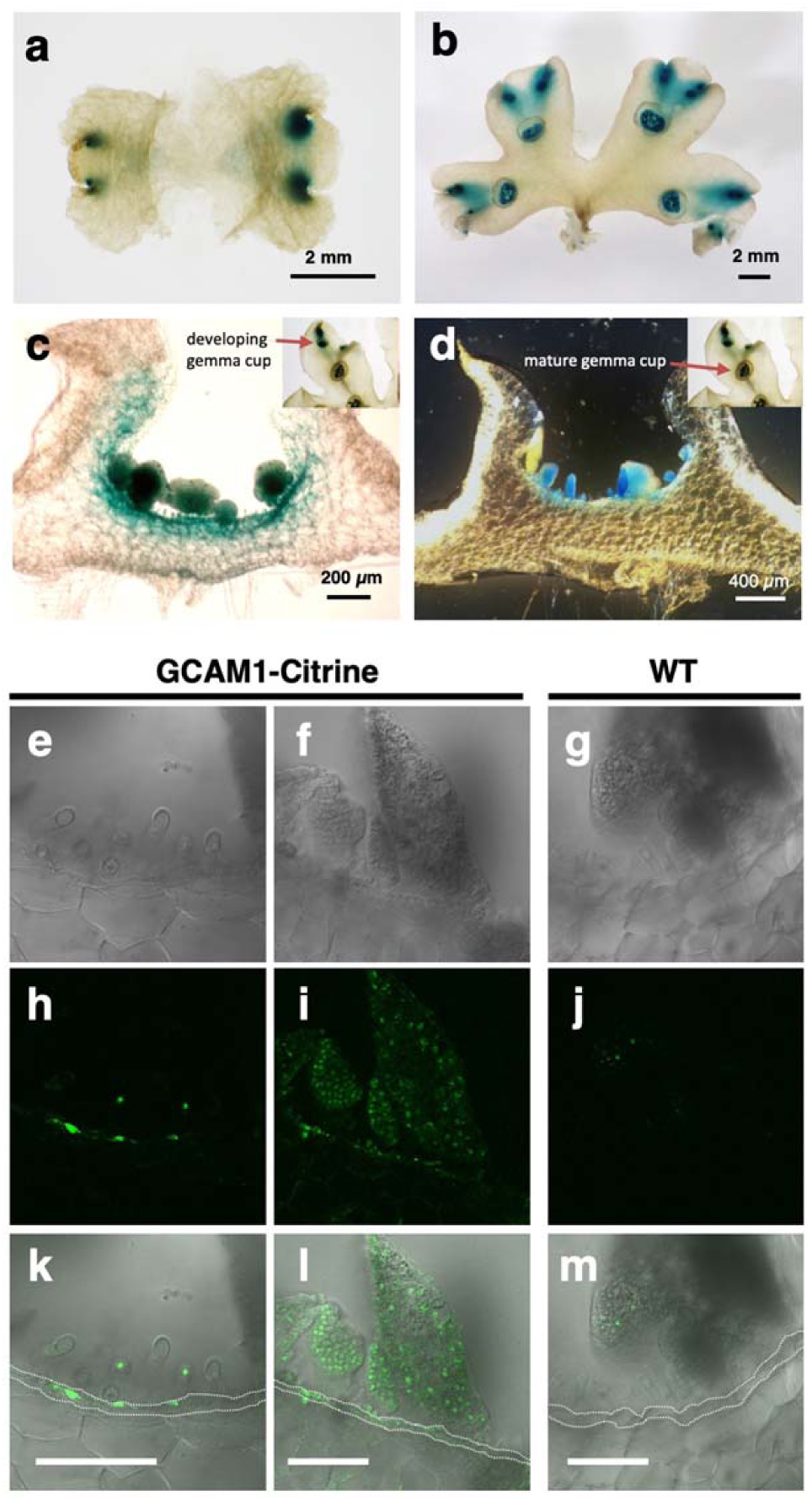
*GCAM1* expression pattern in the vegetative growth stage of *Marchantia polymorpha*. **(a–d)** Histochemical GUS assay in a representative *GCAM1pro:GUS* line. **(a)** Ten-day-old thallus. **(b)** Three-week-old thallus with gemma cups. **(c and d)** Transverse sections of developing and mature gemma cups on a 3-week-old thallus. Developing gemma cups are those closely located near the apical notch **(c)** and mature gemma cups are those located in the basal region of the thallus **(d)**. **(e–m)** Accumulation of GCAM1 in the basal floor of the gemma cup. Bright-field **(e–g)**, single confocal **(h–j),** and merged **(k–m)** images of the basal floor of gemma cups in 17-to 18-day-old thalli in *GCAM1-Citrine* #1 knock-in plants **(e, f, h, i, k, and l)** and the wild type **(g, j, and m)**. Dotted lines indicate basal floor cells. Bars, 100 µm **(e–m).**

To analyze the localization of protein accumulation, we inserted the yellow fluorescent protein (Citrine) sequence into the 3′ end of the *GCAM1* coding sequence (CDS) in the *M. polymorpha* genome (Supplementary Figure 2a, b). The *GCAM1*-*Citrine* knock-in plants exhibited normal organ development, including gemma cups during vegetative growth (Supplementary Figure 2c). Consistent with the promoter–reporter analysis, we detected Citrine signals in the nucleus of cells in the developing gemma and gemma-cup floor cells (Fig. 2e–m).

### GCAM1 is essential for gemma cup formation

To investigate the function of GCAM1, we disrupted *GCAM1* using homologous recombination-mediated gene targeting technology^16^. Two independent *GCAM1* knock-out lines (*gcam1*^*ko*^) were isolated from 522 candidates (Supplementary Figure 2d, e). In the wild-type 3-week-old thallus developed from a thallus explant containing an apical notch, over ten gemma cups filled with gemmae were observed on the midrib of the dorsal surface (Fig. 3a, c, e). In contrast, no gemma cup development was observed on the dorsal surface of the *gcam1*^*ko*^ midrib (Fig. 3b and d), resulting in no gemma generation. No distinct impairment of thallus growth or development of other vegetative organs, i.e. air chamber, rhizoid, and ventral scales, was observed in *gcam1*^*ko*^ thalli compared with those of wild-type thalli (Fig. 3c–f). We also generated *GCAM1* mutants using CRISPR/Cas9-mediated targeted genome editing^30^. Two independent *GCAM1* mutants for each of two independent target sequences of *GCAM1* exhibited absence of gemma cup development, but no other obvious aberrant phenotype was observed during vegetative growth (Supplementary Figure 3), and thus the phenotypes were essentially identical to those of *gcam1*^*ko*^ plants. Altogether, these results demonstrated that *GCAM1* performs a critical role in gemma cup formation.

**Fig. 3.**
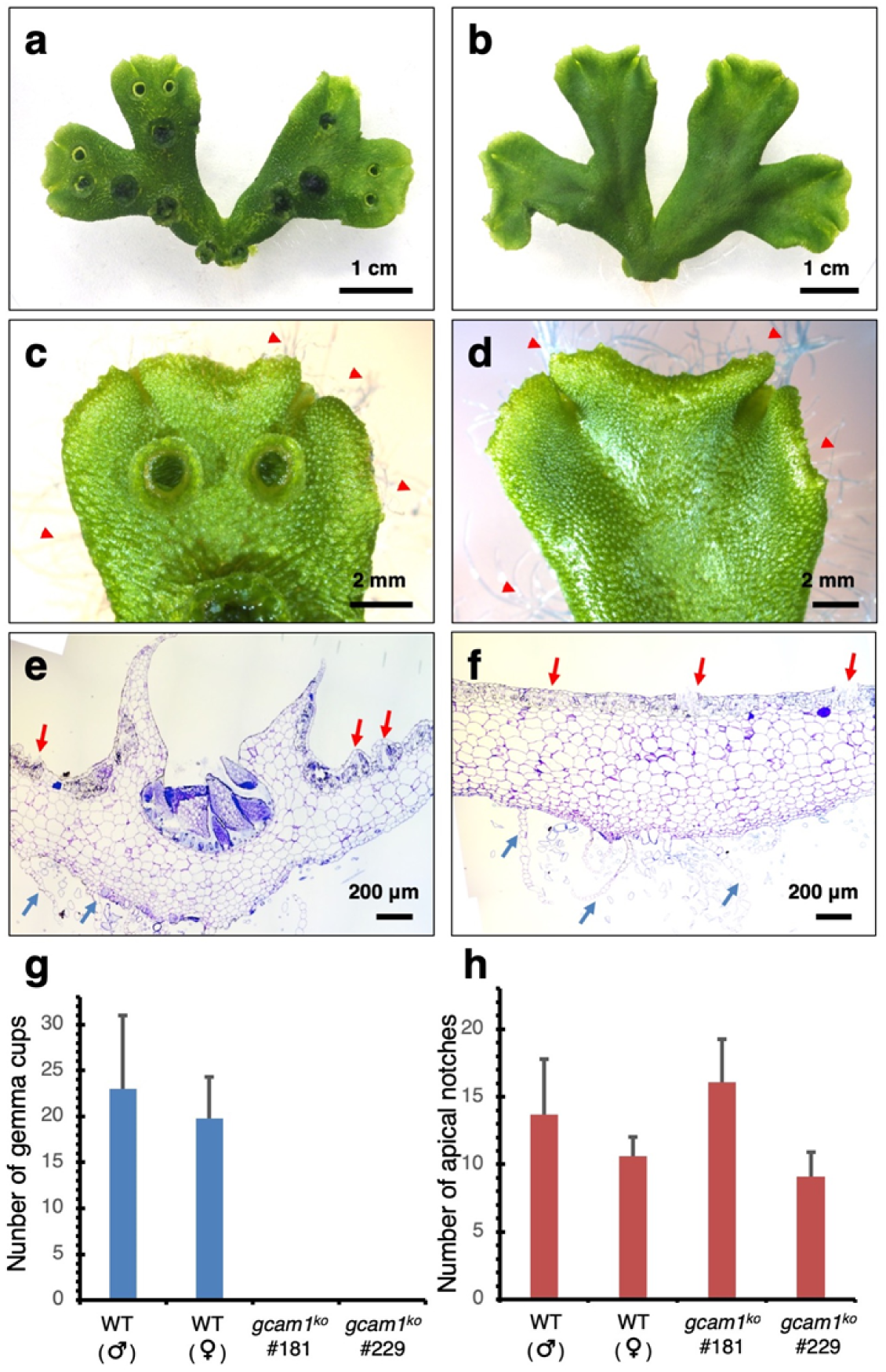
Phenotype of the *gcam1* mutant of *Marchantia polymorpha.* **(a and b)** Three-week-old thalli grown from tip of thalli of the wild type **(a)** and *gcam1*^*ko*^ knock-out mutant **(b)**. The thalli were grown from the tip of parental thalli, because gemmae were absent in *gcam1*^*ko*^ lines. **(c and d)** Close-up of the tip of thalli of the previous individual, showing gemma cups in the wild type **(c)**, whereas gemma cups were absent in the *gcam1*^*ko*^ lines **(d).** Red arrowheads indicate rhizoids. **(e and f)** Transverse sections (5 µm thick) from 3-week-old thalli of the wild type **(e)** and *gcam1*^*ko*^ **(f).** Red and blue arrows indicate air chambers and ventral scales, respectively. **(g)** Number of gemma cups in the wild type (WT) and *gcam1*^*ko*^ lines. **(h)** Number of apical notches in the wild type and *gcam1*^*ko*^ lines. Values are means ± s.d (*n* = 10).

Generation of gemma cups is correlated with bifurcation (i.e. branching of the apical meristem) of thalli^7^. To clarify whether *GCAM1* may also be involved in branching, we investigated the number of gemma cups and apical notches in the *gcam1*^*ko*^ line in comparison with those of the wild type (Fig. 3g, h). About 20 gemma cups were observed on 3-week-old wild-type thalli, whereas no gemma cups developed in the *gcam1*^*ko*^ mutants. In contrast to the conspicuous gemma cup phenotype, no significant decrease in the number of bifurcations was observed in the *gcam1*^*ko*^ lines compared with that of the wild type, which suggested that *GCAM1* plays no role in bifurcation of the apical notch.

### Overexpression of *GCAM1* promotes proliferation of undifferentiated cells

To further investigate the developmental function of GCAM1, we generated transgenic plants that constitutively expressed *GCAM1* under the control of the Mp*ELONGATION FACTOR 1α* regulatory sequence in *M. polymorpha* (Mp*EFpro:GCAM1*). We obtained a significantly lower number of transformants using the Mp*EFpro:GCAM1* construct than usual, and the majority of the obtained Mp*EFpro:GCAM1* lines formed a mass of tiny flat thallus-like structures (Supplementary Figure 4). Thus, we utilized an inducible system (Fig. 4a), in which protein nuclear localization is modulated by a glucocorticoid receptor (GR) domain^31,32^. In these transgenic plants, GCAM1 function was expected to be induced by treatment with dexamethasone (DEX). In the wild type, treatment with DEX caused no morphological effect (Supplementary Figure 5). In the absence of DEX, Mp*EFpro:GCAM1-GR* plants exhibited the wild-type phenotype and formed normal dorsal/ventral organs, i.e. air chambers, gemma cups, scales, and rhizoids (Fig. 4b, d, f). Treatment of Mp*EFpro:GCAM1-GR* plants with DEX severely suppressed thallus growth, and cell clumps with no dorsal/ventral organs and few rhizoids were observed (Fig. 4c, e, and g). After suspension of the DEX treatment, a number of small thallus branches were generated in random positions (Fig. 4h, i and Supplementary Figure 6). These results implied that GCAM1 functions in the maintenance and proliferation of pluripotent cells that possess the capability to develop into individual thalli.

**Fig. 4.**
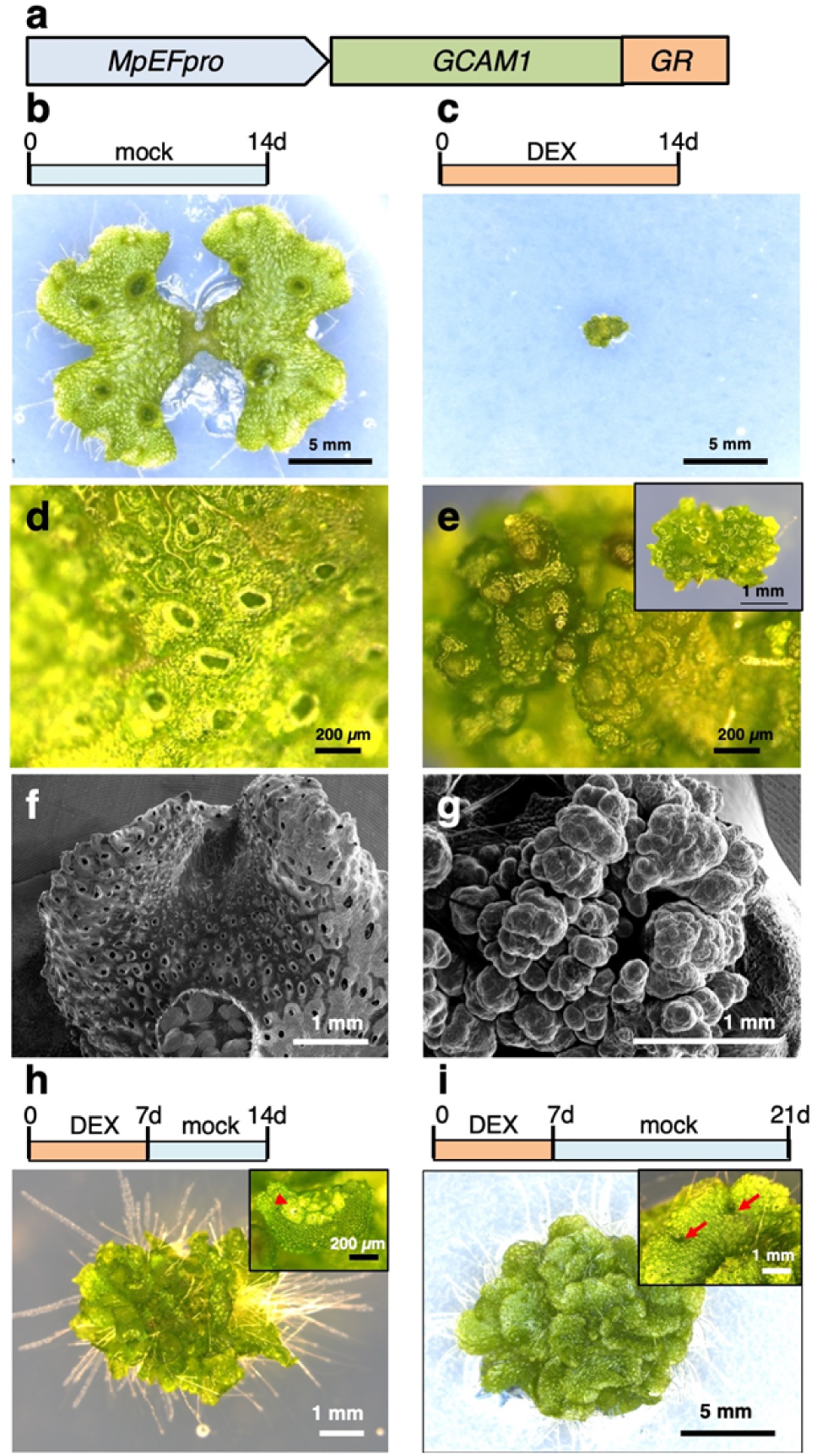
Induction of GCAM1 function suppresses growth and organ development in thallus of *Marchantia polymorpha.* **(a)** Schematic representation of the Mp*EFpro:GCAM1-GR* construct. **(b, d, f)** Two-week-old Mp*EFpro:GCAM1-GR* transgenic plant treated with mock. A number of air pores and air chambers are shown in high-magnification views of the thallus surface **(d and f). (c, e, and g)** Two-week-old Mp*EFpro:GCAM1-GR* transgenic plant treated with 10 µM DEX. There is no indication of air chamber development **(e and g).** Much fewer rhizoids developed from the ventral surface **(e, inset). (h)** Two-week-old Mp*EFpro:GCAM1-GR* transgenic plant treated with DEX for the first 7 days and mock for the latter 7 days, and a high-magnification view of the thallus surface (inset). Red arrowhead indicates a developing air chamber. **(i)** Three-week-old Mp*EFpro:GCAM1-GR* transgenic plant treated with DEX for the first 7 days and mock for the latter 14 days, and a high-magnification view of the thallus (inset). Red arrows indicate a set of bifurcated apical notches of thalli.

### Orthologous relationship of GCAM1 to R2R3-MYBs in subfamily 14

Gemmae and gemma cups are specialized and derived gametophytic organs for vegetative reproduction in certain Marchantiopsida species, a group of complex thalloid liverworts including *M. polymorpha*. However, orthologs of *GCAM1* have been identified across diverse branches of land plants. The inferred amino acid sequence of GCAM1 contained an R2R3-MYB DNA-binding domain towards the N-terminus, and showed the highest similarity (3e-59) to REGULATOR OF AXILLARY MERISTEM 3 (RAX3) in a BLASTP search against the *Arabidopsis thaliana* protein database. Phylogenetic analysis suggested that GCAM1 belonged to a clade, the R2R3-MYB subfamily 14, which includes the Arabidopsis *RAX* genes and tomato *Blind* gene (Fig. 5a), with highly conserved exon– intron structures (Fig. 5b). Some angiosperm proteins in the R2R3-MYB subfamily 14 are regulatory factors for axillary meristem formation, e.g. *Blind* and *RAX* genes^33,34,35,36^. Relative to other R2R3-MYB proteins, those of the R2R3-MYB subfamily 14 are characterized by an additional amino acid insertion between the first and second conserved Trp residues of the R2 repeat^28^, and GCAM1 conforms to this rule (Fig. 5c). These findings support the orthologous relationship of GCAM1 to angiosperm proteins in the R2R3-MYB subfamily 14.

**Fig. 5.**
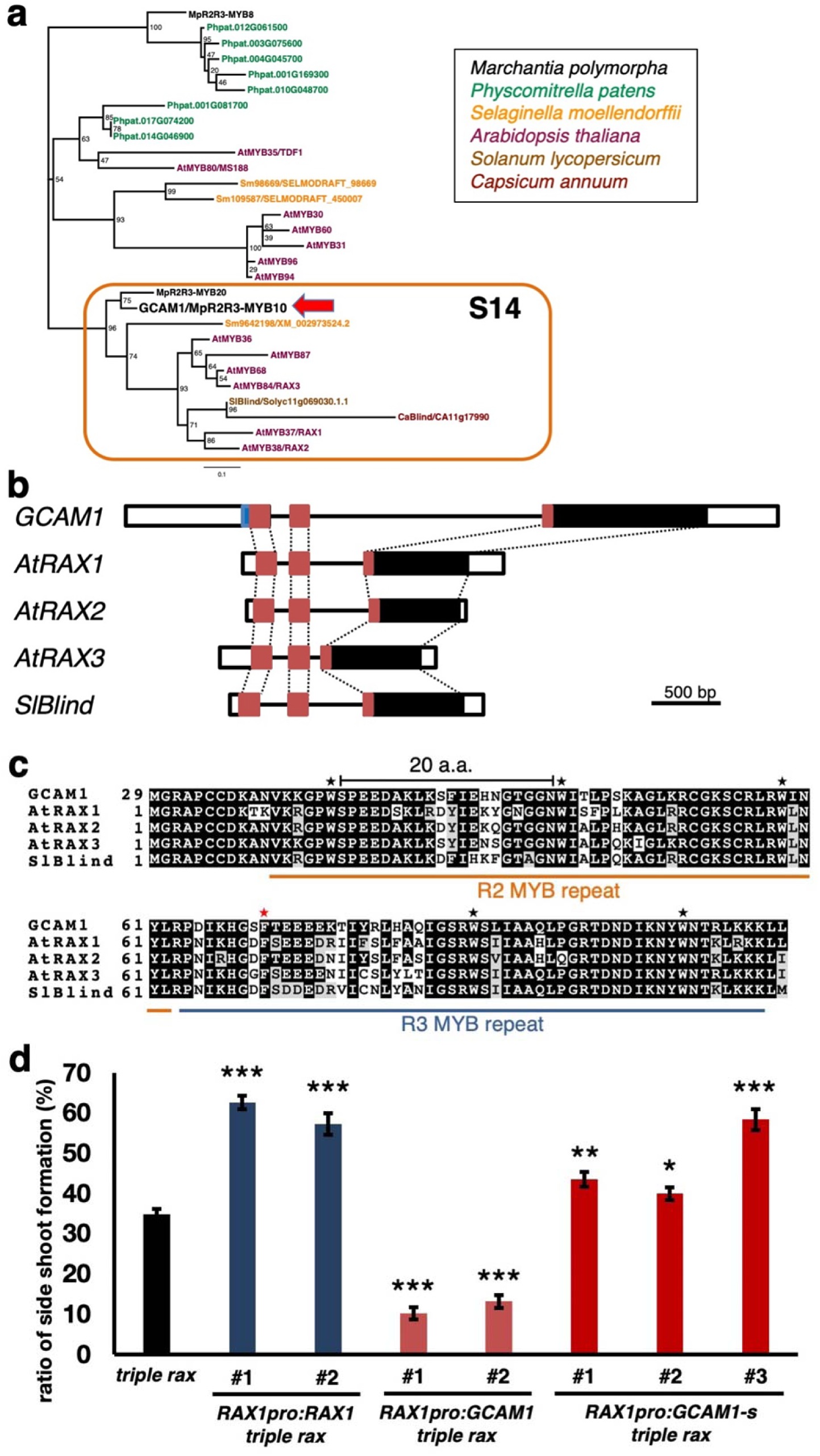
Relationships of GCAM1 and angiosperm R2R3-MYB subfamily 14 orthologs. **(a)** Phylogenetic analysis of *GCAM1* and its homologs in plants. Unrooted maximum-likelihood tree generated from the amino acid sequences for the R2R3-MYB DNA-binding domain of GCAM1 and related R2R3-MYB proteins across diverse plant lineages. The numbers at the nodes are bootstrap values calculated from 1000 replicates. The scale bar is evolutionary distance as the rate of amino acid substitutions. *SlBlind* and *CaBlind* are R2R3-MYB genes reported to function in axillary bud formation in *Solanum lycopersicum* and *Capsicum annuum*, respectively. From *A. thaliana*, not only R2R3-MYBs in subfamily 14, but also other R2R3-MYBs in closely related subfamilies were sampled for the phylogenetic analysis. For *Physcomitrella patens* and *Selaginella moellendorffii*, R2R3-MYBs that showed eight highest and two highest similarities to GCAM1, respectively, were sampled. **(b)** Exon–intron structure of *GCAM1, AtRAX1, AtRAX2, AtRAX3*, and *SlBlind*. Box, exon; line, intron; white, 5′- or 3′-untranslated region. Red and blue boxes indicate R2R3-MYB domains and the non-conserved N-terminal stretch of 28 a.a. in GCAM1, respectively. **(c)** Multiple alignment of R2R3-MYB domains of GCAM1, AtRAX1, AtRAX2, AtRAX3, and SlBlind. Black asterisks indicate conserved Trp residues in plant R2R3-MYBs, and the red asterisk indicates a Phe residue conserved in R2R3-MYB subfamily 14^28^. R2R3-MYBs in subfamily 14 are characterized by an additional amino acid (20 a.a.) compared with other R2R3-MYB subfamilies (19 a.a.) between the first and the second Trp residues of the R2 repeat. **(d)** Genetic complementation of Arabidopsis *rax1-3 rax2-1 rax3-1* mutant with Arabidopsis *RAX1* and Marchantia *GCAM1* (see Supplementary Figure 7 for details). Lateral shoot formation in rosette leaf axils of the *rax1-3 rax2-1 rax3-1* mutant (*triple rax, n* = 30), two independent transformants containing *RAX1*-promoter-driven *RAX1* open reading frame (ORF1) in *rax* triple mutant background (*RAX1pro:RAX1 triple rax, n* = 15), two independent transformants containing *RAX1*-promoter-driven *GCAM1* ORF in *rax* triple mutant background (*RAX1pro:GCAM1 triple rax, n* = 15), and three independent transgenics containing *RAX1*-promoter-driven shorter version of *GCAM1* ORF in *rax* triple mutant background (*RAX1pro:GCAM1-s triple rax, n* = 15). Values represent means ± SE. Asterisks indicate significant differences relative to triple *rax* mutant (* *P* < 0.05, ** *P* < 0.02, *** *P* < 0.0001).

To examine whether *M. polymorpha GCAM1* is functionally equivalent to the angiosperm orthologs and is able to function in control of axillary meristem formation in angiosperms, we introduced *GCAM1* into the Arabidopsis *rax1 rax2 rax3* triple mutant, which exhibits significant inhibition of axillary bud formation compared with the wild type^34^. Interestingly, expression of *GCAM1* under the control of the Arabidopsis *RAX1* promoter (*RAX1pro:GCAM1*) further inhibited axillary meristem formation in the *rax1 rax2 rax3* triple mutant background (Fig. 5 and Supplementary Figure 7). Given that a non-conserved N-terminal domain is present upstream of (28 amino acids from) the R2R3-MYB domain of *GCAM1*, which does not exist in angiosperm R2R3-MYB subfamily 14 members (Fig. 5b, c), we also tested complementation with a shorter open reading frame of *GCAM1* (*GCAM1-s*), which starts from the second start codon in the original *GCAM1* ORF. The *RAX1pro:GCAM1-s* transgenic plants showed notable recovery of axillary meristem formation compared with that of the *rax* triple mutant (Fig. 5 and Supplementary Figure 7). These results demonstrated the capability of *M. polymorpha* GCAM1 to interact with the genetic machinery of axillary meristem formation in the angiosperm *A. thaliana*.

## Discussion

### Identification of an essential regulator for vegetative reproduction in *M. polymorpha*

The basis of vegetative reproduction in seed plants is the *de novo* meristem formation, which allows development of clonal plants. At present, the molecular mechanism of natural vegetative reproduction is poorly known. In the current study, we identified a key regulator for vegetative reproduction in the basal land plant *M. polymorpha*. Using comprehensive transcriptome analysis and molecular genetic approaches, we identified an R2R3-MYB gene, *GCAM1*, which is an essential factor for gemma cup development in *M. polymorpha*. Loss-of-function mutants of *GCAM1* showed no sign of gemma cup development, but no obvious abnormality was observed in the development of other thallus tissues (Fig. 3). This observation suggested that GCAM1 plays a specific role in gemma cup development in the vegetative growth phase of *M. polymorpha*. Our observations on the accumulation of *GCAM1* transcripts in developing gemma cups and gemmae (Fig. 2e–m) corroborated the suggestion of a specific function for *GCAM1* in gemma cup and gemma formation. However, continuous expression of *GCAM1* did not confer excessive production of gemma cups (Fig. 4) and instead suppressed organ differentiation and promoted generation of undifferentiated cell clumps (Fig. 4 and Supplementary Figure 4). After GCAM1 induction was halted at 1 week or 2 weeks after DEX treatment, numerous thalli developed directly from the cell clumps (Fig. 4), suggesting that the cell clumps consist of cells with the potential to differentiate into meristematic apical cells of the thallus. These results suggest that GCAM1 might function to suppress tissue differentiation and maintain the undifferentiated status of cells, which may be essential for the floor cells of the gemma cup to form gemma initials. The role of GCAM1 would be to proliferate floor cells of gemma cups, and to maintain the somewhat undifferentiated status of these cells, which may be a prerequisite for gemmae and gemma cup development.

### A common developmental mechanism between gemma cups and boundary zone for axillary meristems

The present phylogenetic analysis indicated that *GCAM1* is orthologous to angiosperm genes in the R2R3-MYB subfamily 14^28^, which includes regulatory factors for axillary meristem formation, e.g. tomato *Blind* and Arabidopsis *RAX* genes^33,34,35,36^ (Fig. 5a). Interspecific complementation analyses demonstrated that *M. polymorpha* GCAM1 can interact with the genetic machinery of axillary meristem formation in Arabidopsis (Fig. 5d). These results suggested there is a common regulatory mechanism between vegetative reproduction in the gametophyte body of the liverwort *M. polymorpha* and axillary meristem formation in the sporophyte body of angiosperms. In Arabidopsis, RAX1 and RAX3 are specifically expressed in the boundary region, axils, between the shoot apical meristem (SAM) and leaf primordia, and regulate axillary meristem formation during early vegetative development^34,35^. In angiosperms, boundary zones are characterized by a lower rate of cell divisions and lower concentrations of auxin and brassinosteroids than those of leaf primordia, resulting in a low degree of differentiation of boundary cells that exhibit higher competence for meristem formation^2,37^. In Arabidopsis, *CUP-SHAPED COTYLEDON* genes (*CUCs*) are also known to be expressed in boundary zones and required for boundary formation^2^. The tomato gene *Goblet*, which is the tomato orthologue of *CUCs*, promotes ectopic shoot formation at the leaflet boundary, suggesting that the specification of the boundary zone is the common basis of axillary bud formation and vegetative reproduction in angiosperms^38^. The cells of the gemma cup floor in the liverwort *M. polymorpha*, in which the ability to regenerate clonal plantlets is maintained, could be considered to show similar characteristics to cells in the boundary zones of angiosperms. An orthologous relationship and functional equivalence between *GCAM1* in *M. polymorpha* and *RAX/Blind* genes in angiosperms further implies acquisition of a regulatory mechanism for establishment of the boundary zone, which retains the competence to generate meristematic cells, in the common ancestor of land plants, and such a regulatory mechanism was co-opted for axillary bud formation in the course of land-plant evolution.

In many plant species capable of vegetative reproduction, clonal propagules are often produced as modified forms of axillary buds, e.g. rhizomes in asparagus (*Asparagus officinalis* L.) and iris (*Iris germanica* L.), tubers in potato (*Solanum tuberosum* L.), and stolons (runners) in strawberry (*Fragaria* × *ananassa*) and wild mint (*Mentha arvensis* L.). In some *Dioscorea* species, including the air potato *Dioscorea bulbifera*, aerial clonal propagules (bulbils) are produced in the leaf axils and detach from the parental plant. In addition, aerial tubers are often formed in the leaf axils of stressed potato plant^39^. Ectopic expression of the tomato cytokinin-activating enzyme gene LONELY GUY (LOG1) transforms axillary meristems into aerial tubers^40^. Thus, there is a strong link between vegetative reproduction and axillary bud formation in angiosperms that is convergent to the clonal reproduction observed in complex thalloid liverworts. The present findings on the liverwort *M. polymorpha* suggests a homology in the mechanisms controlling vegetative reproduction and axillary bud formation in land plants. Additional molecular studies on the process of vegetative reproduction in diverse lineages, such as basal tracheophytes, other bryophytes, or charophycean algae, should shed light on the validity of an evolutionarily conserved gene regulatory network for vegetative reproduction or axillary bud formation in land plants.

## Materials and Methods

### Plant material and growth conditions

Male and female accessions of *M. polymorpha*, Takaragaike-1 (Tak-1) and Takaragaike-2 (Tak-2), respectively^14^, were maintained asexually. F_1_ spores generated by crossing Tak-2 and Tak-1 were used for generation of *gcam1*^*ko*^ and *GCAM1-Citrine* knock-in plants. Formation of sexual organs was induced by far-red irradiation as described previously^41^. Mature sporangia were collected 3–4 weeks after crossing, air-dried for 7–10 days, and stored at −80°C until use.

*Marchantia polymorpha* plants were cultured using half-strength Gamborg’s B5 medium^42^ containing 1% agar under 50–60 µmol photons m^-2^ s^-1^ continuous white light with a cold cathode fluorescent lamp (OPT-40C-N-L; Optrom, Miyagi, Japan) or light-emitting diode (VGL-1200W; SYNERGYTEC, Tokushima, Japan) at 22°C unless otherwise defined.

### RNA extraction and Illumina sequencing

Developing gemma cups located within 3 mm from an apical notch of 3-week-old thalli were manually dissected and immediately immersed in RNAlater (Thermo Fisher Scientific, Waltham, MA, USA). For the control, the midrib region between gemma cups of 3-week-old thalli and 1-week-old thalli, which had no visible gemma cups, were also sampled. The collected samples were divided into three pools for extraction of total RNA.

Total RNA was extracted with the RNeasy^®^ Plant Mini Kit (QIAGEN) in accordance with the manufacturer’s protocol. The quality and quantity of resultant total RNA were evaluated using a NanoDrop™ 1000 spectrophotometer (Thermo Fisher Scientific), Qubit^®^ 2.0 Fluorometer (Thermo Fisher Scientific), and a Bioanalyzer RNA6000 Nano Kit (Agilent Technologies, Santa Clara, CA, USA). The sequence libraries were constructed with a TruSeq™ RNA Sample Prep Kit v2 (Illumina, San Diego, CA, USA) in accordance with the manufacturer’s protocol. The quality and quantity of each library were determined using a Bioanalyzer with High Sensitivity DNA kit (Agilent Technologies), and KAPA Library Quantification Kit for Illumina (Illumina). Equal amounts of each library were mixed to make the 2 nM pooled library. Illumina sequencing was performed using a HiSeq 2000 platform (Illumina) to produce 101 single-end data. All reads obtained have been deposited in the DDBJ and are available through the Sequence Read Archive (SRA) under the accession number DRA005912.

### RNA-seq data analysis

The resultant reads were mapped to the draft genome of *M. polymorpha* version 3.1 by TopHat ver 2.1.1^43^ with the default parameters. To identify differentially expressed genes (DEGs), the mapped reads from different samples were compared by Cuffdiff 2.2.1^44^ using the default parameters and transcript annotation generated on the Marchantia genome assembly ver. 3.1^12^. Among DEGs, genes with a log2-fold change >1 were considered as preferentially expressed in the gemma cup. The supercomputers of the Academic Center for Computing and Media Studies at Kyoto University and of the Computing Center at the Faculty of Biology-Oriented Science and Technology, Kindai University were used for computation.

### Quantitative RT-PCR analysis

One microgram of total RNA was reverse-transcribed in a 20 μl reaction mixture using ReverTra Ace^®^ qPCR RT Master Mix with gDNA remover (TOYOBO). After the reaction, the cDNA samples were diluted with 220 µl distilled water and 2 μl aliquots were amplified with the LightCycler^®^ Nano Real-time PCR Detection System (Roche Applied Science, Tokyo, Japan) using the KOD SYBR^®^ qRT-PCR Mix (TOYOBO). The PCR cycling program was performed in accordance with the manufacturer’s protocol. The primers used in these experiments are listed in Supplementary Table 3. Mp*EF1α* was used as an internal control.

### Generation of transformants for promoter–reporter analysis

To construct *GCAM1pro:GUS*, the *GCAM1* genomic region, including a 5593-bp fragment upstream of the 83rd Arg codon in the second exon, was amplified from Tak-1 genomic DNA by PCR with the primers GCAM1pro_L1 and GCAM1pro_R2 and cloned into the pENTR™/D-TOPO^®^ cloning vector (Life Technologies, Rockville, MD, USA). This entry vector was used in the Gateway^®^ LR reaction (Life Technologies) with the Gateway binary vector pMpGWB104^45^ to generate the *GCAM1pro:GUS* binary construct, in which the *GUS* reporter gene was translationally fused at the middle of the second exon of *GCAM1*. The *GCAM1pro:GUS* vectors were introduced into regenerating thalli of Tak-1 by *Agrobacterium*-mediated transformation as previously described^15^. Transformants were selected with 10 µg/ml hygromycin B and 100 µg/ml cefotaxime.

### Generation of *gcam1*^*ko*^ and *GCAM1-Citrine* knock-in plants

For knock-in and knock-out experiments, Tak-1 genomic sequences were amplified by PCR with KOD FX Neo DNA polymerase (TOYOBO) and the primer pair listed in Supplementary Table 3. The amplified products were inserted into the *Pac*I and *Asc*I sites of pJHY-TMp^16^ and pJHY-TMp1-Cit^46^ to generate *gcam1*^*ko*^ and *GCAM1-Citrine* transformants, respectively. The vectors were introduced into germinating spores via *Agrobacterium tumefaciens* GV2260 as described previously^16^. Transformants were selected with 10 µg/ml hygromycin B and 100 µg/ml cefotaxime. The transformants harboring the targeted insertions were selected by PCR with KOD FX Neo DNA polymerase and the primer pairs listed in Supplementary Table 3 and Supplementary Figure 2.

### CRISPR/Cas9-based genome editing of *GCAM1*

Loss-of-function mutants of *GCAM1* were generated with the CRISPR/Cas9 system as described previously^30,47^. We selected two target sequences, one located in the first exon, and the other located in the second exon of *GCAM1* (Supplementary Figure 4). An off-target search was performed using casOT^48^. Synthetic oligo DNAs for respective target sites shown in Supplementary Table 3 were annealed, inserted into the entry vector pMpGE_En01, and then introduced into the destination vector pMpGE011^30,47^. The vectors were introduced into regenerating thalli of Tak-1 via *A. tumefaciens* GV2260^15^, and transformants were selected with 0.5 µM chlorsulfuron. Genomic DNA was isolated from transformants and amplified from the target region by PCR. The PCR product was used for sequencing of the respective target sites with an ABI 3100 genetic analyzer (Applied Biosystems, Foster City, CA, USA).

### Visualization of GCAM1-Citrine

Transverse sections of *GCAM1-Citrine* knock-in thalli with apical notches were generated with a pair of razors, and observed under a confocal laser scanning microscope (Olympus FLUOVIEW FV1000, Tokyo, Japan). Excitation and emission wavelengths for Citrine fluorescence were 515 nm (Ar laser) and 520–570 nm, respectively.

### Histology and light microscopy

For histochemical GUS staining, *GCAM1pro:GUS* plants were grown on half-strength B5 medium for respective periods under continuous white light. GUS staining was performed as described previously and at least five independent lines were observed for GUS staining patterns using S8APO (Leica Microsystems) and DMS1000 (Leica) stereoscopic microscopes, or an upright light microscope Axio Scope.A1 (Zeiss) equipped with an AxioCam ERc5c camera (Zeiss).

For plastic-embedded sectioning, 3-week-old thalli developed from gemmae were dissected into small pieces and transferred to fixative solution (2% glutaraldehyde in 0.05 M phosphate buffer, pH 7.0), evacuated with a water aspirator until the specimens sank, and fixed for 2 days at room temperature. The samples were dehydrated in a graded ethanol series and embedded in Technovit 7100 plastic resin. Semi-thin sections (5 μm thickness) for light microscopy were obtained with a microtome (HM 335E, Leica Microsystems, Heerbrugg, Switzerland) and stained with toluidine blue O.

For scanning electron microscopy, plant samples were frozen in liquid nitrogen and directly observed on a VHX-D500 microscope (KEYENCE, Osaka, Japan).

### Generation of *GCAM1*-overexpressing plants

To generate Mp*EFpro:GCAM1*, the *GCAM1* coding sequence was amplified by PCR using KOD Plus Neo DNA polymerase (TOYOBO) with the primer set GCAM1-cds-L1/GCAM1-cds-sR and cloned into the pENTR/D-TOPO cloning vector. The entry vector was used in the Gateway LR reaction with the Gateway binary vector pMpGWB103^45^. The *MpEFpro:GCAM1* vector was introduced into regenerating thalli of Tak-1 as previously described^15^. Transformants were selected with 10 µg/ml hygromycin B and 100 µg/ml cefotaxime.

To construct Mp*EFpro:GCAM1-GR*, the *GCAM1* CDS was amplified by PCR using KOD Plus Neo DNA polymerase (TOYOBO) with the primer set GCAM1-cds-L1/GCAM1-cds-nsR and cloned into the pENTR/D-TOPO cloning vector. The entry vector was used in the Gateway LR reaction with the Gateway binary vector pMpGWB313^45^. The Mp*EFpro:GCAM1-GR* vector was introduced into regenerating thalli of Tak-1 as previously described^15^. Transformants were selected with 0.5 µM chlorsulfuron and 100 µg/ml cefotaxime.

The transformants and mutants were named consistent with the nomenclatural rules for *M. polymorpha*^29^.

### Phenotypic analysis of Mp*EFpro:GCAM1-GR* plants

In the vegetative phase, DEX treatment of *MpEFpro:GCAM1-GR* plants was performed by culture in half-strength Gamborg’s B5 medium containing 10 µM DEX or applying 200 µL of 10 µM DEX solution to plants.

### Phylogenetic analysis of GCAM1

For phylogenetic analysis of GCAM1 and homologous R2R3-MYBs, sequence information was obtained from the PlantTFDB^49^ and Phytozome databases. A multiple alignment of amino acid sequences of GCAM1 and homologous R2R3-MYBs was first constructed using COBALT^50^ and then using the MUSCLE program^51^ implemented in SeaView version 4.5.4^52^ with the default parameters. The alignment is available as Supplemental Data Set 4. The conserved region covering the R2R3-MYB domain was extracted before phylogenetic tree construction, which was performed using the maximum likelihood method with PhyML^53^ with the LG substitution model. A bootstrap analysis with 1000 replicates was performed in each analysis to assess statistical support for the topology. The phylogenetic tree was visualized using FigTree (http://tree.bio.ed.ac.uk/software/figtree/).

### Introduction of *GCAM1* into an Arabidopsis *rax* triple mutant

To generate a RAX1 promoter construct, the upstream region of Arabidopsis *RAX1* including the promoter and 5′ untranslated region (2939 bp) were amplified by PCR using the primers Myb37-pro-SbfIF and Myb37-pro-AscIR. The resulting PCR fragment was inserted into the *Sbf*I and *Asc*I sites of the pGPTV-BAR-AscI vector^54^ in front of the *GUS* gene (Vector pFY124). A GFP fragment was amplified using the primers GFPAscIF and GFPpolylinkerR. This PCR fragment was introduced into the *Asc*I and *Sac*I sites of pFY124 introducing a new *Avr*II site for the following constructions (Vector pQW154).

The *RAX1* open reading frame was amplified by PCR using the primers RAX1AscIF and RAX1SacIR. The PCR fragment was inserted into the *Asc*I and *Sac*I sites of pQW154. The GCMA1 open reading frame was amplified using the primers GCAM1AscIF and GCAM1AvrIIR. The PCR fragment was inserted into the *Asc*I and *Avr*II sites of pQW154. The short open reading frame of the GCAM1 coding sequence (GCAM1-s;from +85 to the stop codon) was amplified using the primers GCAM1NewAscIF and GCAM1AvrIIR. The PCR fragment was inserted into the *Asc*I and *Avr*II sites of pQW154.

The RAX1pro:RAX1, RAX1pro:GCAM1, and RAX1pro:GCAM1-s constructs were introduced into the Arabidopsis *rax1-3 rax2-1 rax3-1* mutant^34^ using the floral dip method, and T_3_ homozygous transgenic lines were used for further analysis.

For cultivation under short-day conditions, Arabidopsis plants were grown in a controlled environment room under a 8 h/16 h (light/dark) photoperiod, 20°C/18°C (day/night) temperatures, and 50% relative humidity. Flowering was induced after 4 weeks by transferring the plants to a 16 h/8 h (light/dark) photoperiod. Cultivation under long-day conditions was done in a controlled greenhouse with additional artificial light when needed.

Axillary bud formation was analyzed after the onset of flowering using a stereomicroscope as previously described^55^. Buds that had produced one or two leaf primordia as well as elongating lateral shoots were scored as leaf axils that had established axillary meristems.

## Supporting information

A combined file for all supplemental figures and tables

## Data Availability

All reads obtained have been deposited in the DDBJ and are available through the Sequence Read Archive (SRA) under the accession number DRA005912. The authors declare that all data supporting the findings of this study are available within the manuscript and its supplementary files or are from the corresponding author upon request.

## Supplemental Data

The following materials are available in the online version of this article.

**Supplementary Figure 1.** Histological description for the origin and development of gemma cup in *Marchantia polymorpha*.

**Supplementary Figure 2.** Generation of *GCAM1-Citrine* knock-in and *gcam1*^*ko*^ plants, related to Figs. 2 and 3.

**Supplementary Figure 3.** Targeted disruption of *GCAM1* using CRISPR/Cas9-mediated genome editing, related to Fig. 3, related to Fig. 3.

**Supplementary Figure 4.** Overexpression of *GCAM1*, related to Fig. 4.

**Supplementary Figure 5.** No morphological phenotype is conferred by DEX treatment in wild-type thalli, related to Fig. 4.

**Supplementary Figure 6.** Induction of GCAM1 function suppresses growth and organ development, related to Fig. 4.

**Supplementary Figure 7.** Interaction of GCAM1 with Arabidopsis mechanism for axillary meristem formation, related to Fig. 5.

**Supplementary Table 1.** Summary of the RNA-sequencing analysis.

**Supplementary Table 2.** Transcription factor genes upregulated in the gemma cup.

**Supplementary Table 3.** Primers used in this study.

**Supplemental Data Set 1.** List of 1391 genes upregulated by >2 times in the gemma cup in comparison with the young thallus (GC vs TH in Fig. 1b)

**Supplemental Data Set 2.** List of 1483 genes upregulated by >2 times in the gemma cup in comparison with the midrib (GC vs MR in Fig. 1b)

**Supplemental Data Set 3.** List of 664 genes upregulated by >2 times in the gemma cup in comparison with the young thallus and midrib (overlap of GC vs TH and GC vs MR in Fig. 1b)

**Supplemental Data Set 4.** Multiple alignment of GCAM1 and related R2R3-MYBs used for phylogenetic analysis in Fig. 5a.

## ACKNOWLEDGMENTS

The authors thank John L. Bowman and Eduardo Flores-Sandoval (Monash University, Australia) for critical reading of the manuscript; Andrea Busch, Sabine Zachgo (University of Osnabrück, Germany), Shohei Yamaoka, and members of T. Kohchi’s laboratory for technical assistance and discussions. The authors thank Ursula Pfordt and Alexandra Kalde for excellent technical assistance. Research in the Theres laboratory was supported by the Max Planck Society. The RNA-seq analysis was supported by the NIBB Collaborative Research Program (15-823 to K.I.). This study was supported by MEXT KAKENHI Grants-in-Aid for Scientific Research on Innovative Area (18H04836 to R.N., 25113009 to T.K., and 25119711, 15H01233 and 17H06472 to K.I.), JSPS KAKENHI Grants-in-Aid for Scientific Research (B) (15H04391 to K.I.), and by the SUNTORY Foundation for Life Sciences, the Yamada Science Foundation, and the Asahi Glass Foundation (to K.I.). We thank Robert McKenzie, PhD, from Edanz Group (www.edanzediting.com/ac), for editing a draft of this manuscript.

## AUTHOR CONTRIBUTIONS

K.I., S.T., and Y.Y. designed the research; S.T., Y.Y., T.S., R.N., Q.W., H.K., K.T., and K.I. performed research; Y.Y., S.T., R.N., Q.W., K.T.Y., H.K., K.T., H.F., T.M., T.K., and K.I. analyzed the data; K.I. and Y.Y. wrote the paper.

## REFERENCES

1. Steeves TA, Sussex IM. Patterns in plant development, 2 edn. Cambridge University Press (1989).

2. Wang Q, Hasson A, Rossmann S, Theres K. Divide et impera: boundaries shape the plant body and initiate new meristems. New Phytol, (2015).

3. Steward FC, Mapes MO, Kent AE, Holsten RD. Growth and development of cultured plant cells. Science 143, 20–27 (1964).

4. Steward FC, Mapes MO, Mears K. Growth and organized development of cultured cells. II. Organization in cultures grown from freely suspended cells. Am J Bot 45, 705–708 (1958).

5. Klimešová J, Klimeš L. Bud banks and their role in vegetative regeneration – A literature review and proposal for simple classification and assessment. Perspect Plant Ecol Evol Syst 8, 115–129 (2007).

6. Davis FT, Geneve, R.L., Wilson, S.E., Hartmann, H.T., Kester, D.E. Hartmann & Kester’s Plant Propagation: Principles and Practices. Pearson (2017).

7. Shimamura M. Marchantia polymorpha: Taxonomy, phylogeny and morphology of a model system. Plant Cell Physiol 57, 230–256 (2016).

8. Barnes CR, Land WJG. Bryological papers. II. The origin of the cupule of Marchantia. Bot Gaz 46, 401–409 (1908).

9. Voth PD, Hamner KC. Responses of *Marchantia polymorpha* to nutrient dupply and p dhotoperiod. Bot Gaz 102, 169–205 (1940).

10. Voth PD. Gemmae-cup production in *Marchantia polymorpha* and its response to calcium deficiency and supply of other nutrients. Bot Gaz 103, 310–325 (1941).

11. Benson-Evans K. Physiology of the reproduction of bryophytes. The Bryologist 67, 431–445 (1964).

12. Bowman JL, et al. Insights into land plant evolution garnered from the *Marchantia polymorpha* genome. Cell 171, 287–304 e215 (2017).

13. Ishizaki K, Nishihama R, Yamato KT, Kohchi T. Molecular genetic tools and techniques for *Marchantia polymorpha* research. Plant Cell Physiol 57, 262–270 (2016).

14. Ishizaki K, Chiyoda S, Yamato KT, Kohchi T. *Agrobacterium*-mediated transformation of the haploid liverwort *Marchantia polymorpha* L., an emerging model for plant biology. Plant Cell Physiol 49, 1084–1091 (2008).

15. Kubota A, Ishizaki K, Hosaka M, Kohchi T. Efficient *Agrobacterium*-mediated transformation of the liverwort *Marchantia polymorpha* using regenerating thalli. Biosci Biotechnol Biochem 77, 167–172 (2013).

16. Ishizaki K, Johzuka-Hisatomi Y, Ishida S, Iida S, Kohchi T. Homologous recombination-mediated gene targeting in the liverwort *Marchantia polymorpha* L. Sci Rep 3, 1532 (2013).

17. Flores-Sandoval E, Dierschke T, Fisher TJ, Bowman JL. Efficient and inducible use of artificial microRNAs in *Marchantia polymorpha*. Plant Cell Physiol 57, 281–290 (2016).

18. Nishihama R, Ishida S, Urawa H, Kamei Y, Kohchi T. Conditional gene expression/deletion systems for *Marchantia polymorpha* using its own heat-shock promoter and Cre/loxP-mediated site-specific recombination. Plant Cell Physiol 57, 271–280 (2016).

19. Sugano SS, et al. CRISPR/Cas9-mediated targeted mutagenesis in the liverwort *Marchantia polymorpha* L. Plant Cell Physiol 55, 475–481 (2014).

20. Menand B, et al. An ancient mechanism controls the development of cells with a rooting function in land plants. Science 316, 1477–1480 (2007).

21. Kubota A, Kita S, Ishizaki K, Nishihama R, Yamato KT, Kohchi T. Co-option of a photoperiodic growth-phase transition system during land plant evolution. Nat Commun 5, 3668 (2014).

22. Xu B, et al. Contribution of NAC transcription factors to plant adaptation to land. Science 343, 1505–1508 (2014).

23. Kato H, et al. Auxin-mediated transcriptional system with a minimal set of components is critical for morphogenesis through the life cycle in *Marchantia polymorpha*. PLoS Genet 11, e1005084 (2015).

24. Flores-Sandoval E, Eklund DM, Bowman JL. A simple auxin transcriptional response system regulates multiple morphogenetic processes in the liverwort *Marchantia polymorpha*. PLoS Genet 11, e1005207 (2015).

25. Proust H, et al. RSL class I genes controlled the development of epidermal structures in the common ancestor of land plants. Curr Biol 26, 93–99 (2016).

26. Ishizaki K. Evolution of land plants: insights from molecular studies on basal lineages. Biosci Biotechnol Biochem 81, 73–80 (2017).

27. Pires ND, Dolan L. Morphological evolution in land plants: new designs with old genes. Philos Trans R Soc Lond B Biol Sci 367, 508–518 (2012).

28. Stracke R, Werber M, Weisshaar B. The R2R3-MYB gene family in *Arabidopsis thaliana*. Curr Opin Plant Biol 4, 447–456 (2001).

29. Bowman JL, et al. The Naming of Names: Guidelines for Gene Nomenclature in Marchantia. Plant Cell Physiol 57, 257–261 (2016).

30. Sugano SS, et al. Efficient CRISPR/Cas9-based genome editing and its application to conditional genetic analysis in Marchantia polymorpha. PLoS One 13, e0205117 (2018).

31. Lloyd AM, Schena M, Walbot V, Davis RW. Epidermal cell fate determination in Arabidopsis: patterns defined by a steroid-inducible regulator. Science 266, 436–439 (1994).

32. Schena M, Lloyd AM, Davis RW. A steroid-inducible gene expression system for plant cells. Proc Natl Acad Sci U S A 88, 10421–10425 (1991).

33. Schmitz G, Tillmann E, Carriero F, Fiore C, Cellini F, Theres K. The tomato Blind gene encodes a MYB transcription factor that controls the formation of lateral meristems. Proc Natl Acad Sci U S A 99, 1064–1069 (2002).

34. Müller D, Schmitz G, Theres K. Blind homologous R2R3 Myb genes control the pattern of lateral meristem initiation in Arabidopsis. Plant Cell 18, 586–597 (2006).

35. Keller T, Abbott J, Moritz T, Doerner P. Arabidopsis REGULATOR OF AXILLARY MERISTEMS1 controls a leaf axil stem cell niche and modulates vegetative development. Plant Cell 18, 598–611 (2006).

36. Jeifetz D, David-Schwartz R, Borovsky Y, Paran I. CaBLIND regulates axillary meristem initiation and transition to flowering in pepper. Planta 234, 1227–1236 (2011).

37. Wang Q, Kohlen W, Rossmann S, Vernoux T, Theres K. Auxin depletion from the leaf axil conditions competence for axillary meristem formation in Arabidopsis and tomato. Plant Cell, (2014).

38. Rossmann S, Kohlen W, Hasson A, Theres K. Lateral suppressor and Goblet act in hierarchical order to regulate ectopic meristem formation at the base of tomato leaflets. Plant J 81, 837–848 (2015).

39. Ewing EE, Struik PC. Tuber Formation in potato: induction, initiation, and growth. In: Horticultural Reviews (eds Janick J) John Wiley & Sons, Inc. (1992).

40. Eviatar-Ribak T, Shalit-Kaneh A, Chappell-Maor L, Amsellem Z, Eshed Y, Lifschitz E. A cytokinin-activating enzyme promotes tuber formation in tomato. Curr Biol 23, 1057–1064 (2013).

41. Chiyoda S, Ishizaki K, Kataoka H, Yamato KT, Kohchi T. Direct transformation of the liverwort *Marchantia polymorpha* L. by particle bombardment using immature thalli developing from spores. Plant Cell Rep 27, 1467–1473 (2008).

42. Gamborg OL, Miller RA, Ojima K. Nutrient requirements of suspension cultures of soybean root cells. Exp Cell Res 50, 151–158 (1968).

43. Kim D, Pertea G, Trapnell C, Pimentel H, Kelley R, Salzberg SL. TopHat2: accurate alignment of transcriptomes in the presence of insertions, deletions and gene fusions. Genome Biol 14, R36 (2013).

44. Trapnell C, et al. Differential gene and transcript expression analysis of RNA-seq experiments with TopHat and Cufflinks. Nat Protoc 7, 562–578 (2012).

45. Ishizaki K, et al. Development of Gateway binary vector series with four different selection markers for the liverwort *Marchantia polymorpha*. PLoS One 10, e0138876 (2015).

46. Yamaoka S, et al. Generative cell specification requires transcription factors evolutionarily conserved in land plants. Curr Biol 28, 479–486 e475 (2018).

47. Sugano SS, Nishihama R, CRISPR/Cas9-based genome editing of transcription factor genes in *Marchantia polymorpha*. Methods Mol Biol 1830: 109–126 (2018).

48. Xiao A, et al. CasOT: a genome-wide Cas9/gRNA off-target searching tool. Bioinformatics 30, 1180–1182 (2014).

49. Jin J, et al. PlantTFDB 4.0: toward a central hub for transcription factors and regulatory interactions in plants. Nucleic Acids Res 45, D1040–D1045 (2017).

50. Papadopoulos JS, Agarwala R. COBALT: constraint-based alignment tool for multiple protein sequences. Bioinformatics 23, 1073–1079 (2007).

51. Edgar RC. MUSCLE: multiple sequence alignment with high accuracy and high throughput. Nucleic Acids Res 32, 1792–1797 (2004).

52. Gouy M, Guindon S, Gascuel O. SeaView version 4: A multiplatform graphical user interface for sequence alignment and phylogenetic tree building. Mol Biol Evol 27, 221–224 (2010).

53. Guindon S, Dufayard JF, Lefort V, Anisimova M, Hordijk W, Gascuel O. New algorithms and methods to estimate maximum-likelihood phylogenies: assessing the performance of PhyML 3.0. Syst Biol 59, 307–321 (2010).

54. Überlacker B, Werr W. Vectors with rare-cutter restriction enzyme sites for expression of open reading frames in transgenic plants. Mol Breed 2, 293–295 (1996).

55. Raman S, Greb T, Peaucelle A, Blein T, Laufs P, Theres K. Interplay of miR164, CUP-SHAPED COTYLEDON genes and LATERAL SUPPRESSOR controls axillary meristem formation in *Arabidopsis thaliana*. Plant J 55, 65–76 (2008).

