## Supplementary material for "An evolutionarily conserved mechanism for production of secondary meristems in land plants": A combined file for all supplemental figures and tables

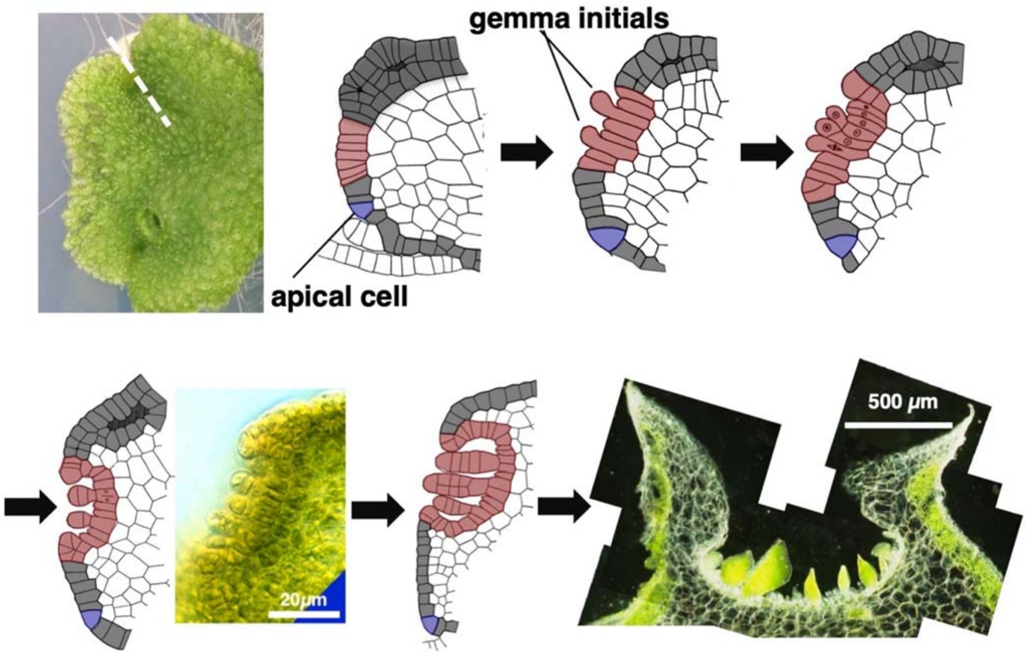


**Supplementary Figure 1. Histological description for the origin and development of gemma cup in *Marchantia polymorpha*.**

Schematic representation of the early stages of gemma cup development. Illustrations of thallus sections are adapted from Barns and Land (1908), and the precursor of gemma cup and gemmae are highlighted in red. Apical cells are indicated in blue. The precursor of gemma cup formation can be recognized as close as third cell back from the apical cell, where a periclinal cell division for air chamber formation is suppressed and continuous single epidermis is observed. The single-layerd epidermal cells will become the basal epidermis of gemma cup floor, and undergo repeated anticlinal cell divisions to enlarge the area of gemma cup floor. Some of gemma cup floor cells begin to develop gemmae from the early stage of gemma cup development, and growth of gema cup occur simultaneously with gemma development and maturation.

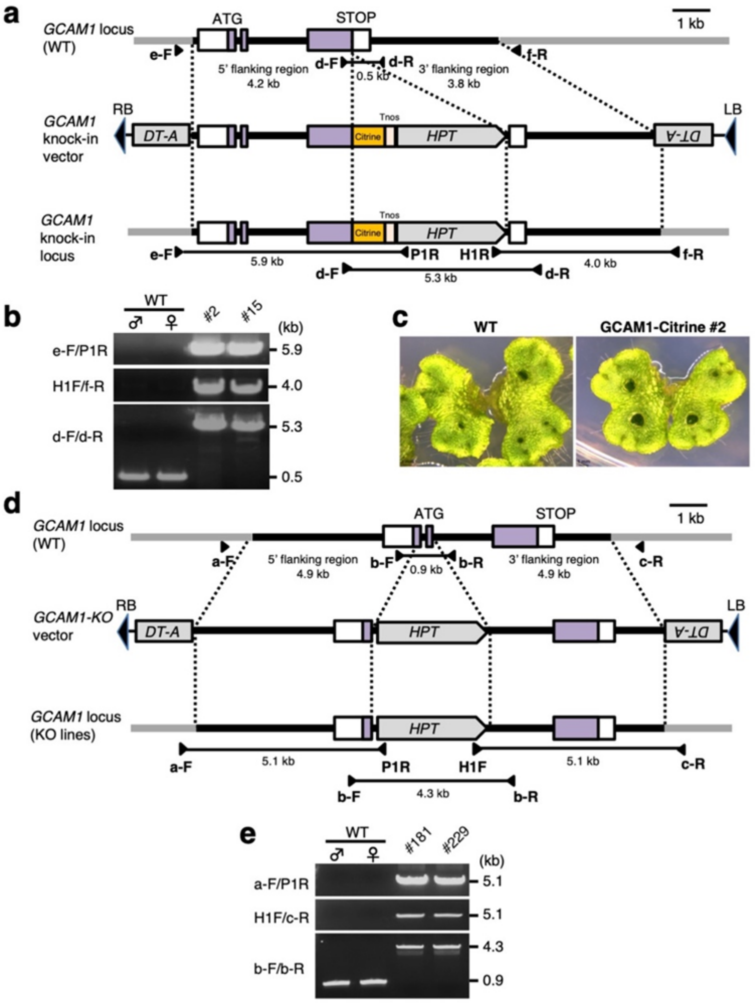


**Supplementary Figure 2. Generation of *GCAM1-Citrine* knock-in and *gcam1*$^{KO}$ plants, related to Fig. 2 and 3.**

**(a)** Structures of *GCAM1* loci in WT and *GCAM1-Citrine* knock-in plants, and the knock-in vector sequences between left (LB) and right (RB) borders. Black bars indicate the flanking sequences subcloned into pJHY-TMp1-Cit. Closed arrowheads indicate primer positions used in (**b**). **(b)** Genomic PCR analysis of *GCAM1* locus in WT and *GCAM1-Citrine* knock-in plants. (**c**) Mature thalli of WT and *GCAM1-Citrine* knock-in plants. There is no morphological abnormality in *GCAM1-Citrine* knock-in plants. (**d**) Structures of *GCAM1* loci in WT and *gcam1*$^{KO}$ and the knockout vector sequences between LB and RB. Black bars indicate the flanking sequences subcloned into pJHY-TMp1, and closed arrowheads indicate primer positions used in (**e**), respectively. (**e**) Genomic PCR analysis of *GCAM1* locus in WT and *gcam1*$^{KO}$ plants. DT-A, diphtheria toxin gene cassette; HPT, hygromycin phosphotransferase gene cassette; Tnos, nopaline synthase gene terminator sequence. White boxes indicate UTRs, and purple boxes indicates protein coding regions.

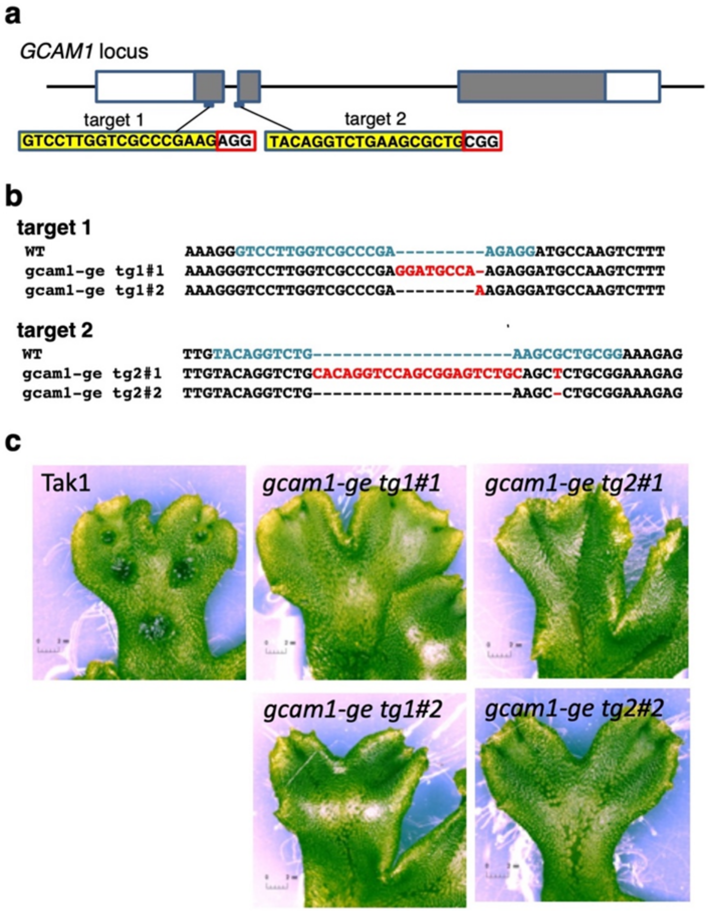


**Supplementary Figure 3. Targeted disruption of *GCAM1* using CRISPR/Cas9-mediated genome editing, related to Fig. 3.**

**(a)** Schematic representation of two independent target sites in *GCAM1*. **(b)** Schematic representation of the construct for CRISPR/Cas9-mediated genome editing in *M. polymorpha*. **(c)** Genome sequences of indicated target sites in wild type and targeted transgenics. (**d)** Top view of 2-week-old thallus grown from tip of thalli in wild type and indicated transgenics.

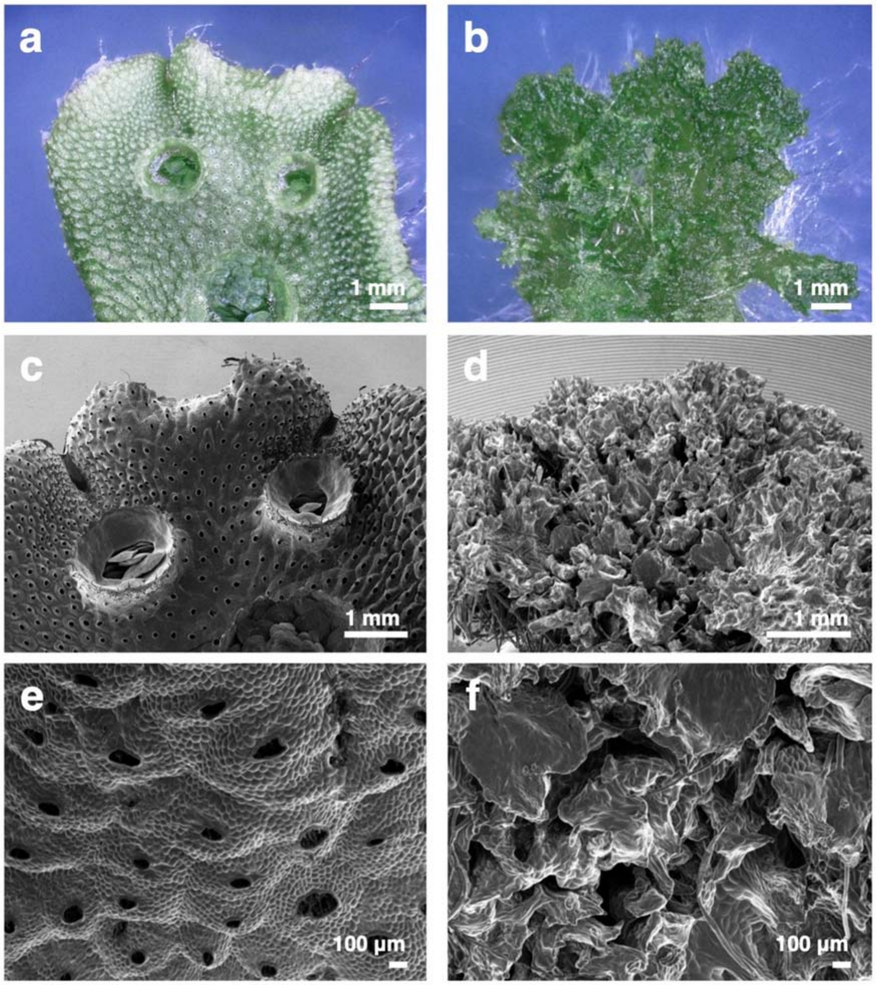


**Supplementary Figure 4. Overexpression of *GCAM1*, related to Fig. 4.**

**(a and b)** 2-week-old thalli grown from tip of thalli of the wild type **(a)** and Mp*EFpro::GCAM1* **(b)** thalli. **(c to f)** Scanning electron micrographs of wild-type **(c, e)** and Mp*EFpro::GCAM1* **(d, f)** thalli. Close-up of dorsal surface of previous individual, showing numerous air pores in the wild type **(e)**, whereas no air pores were observed in the Mp*EFpro::GCAM1* **(f)** thalli.

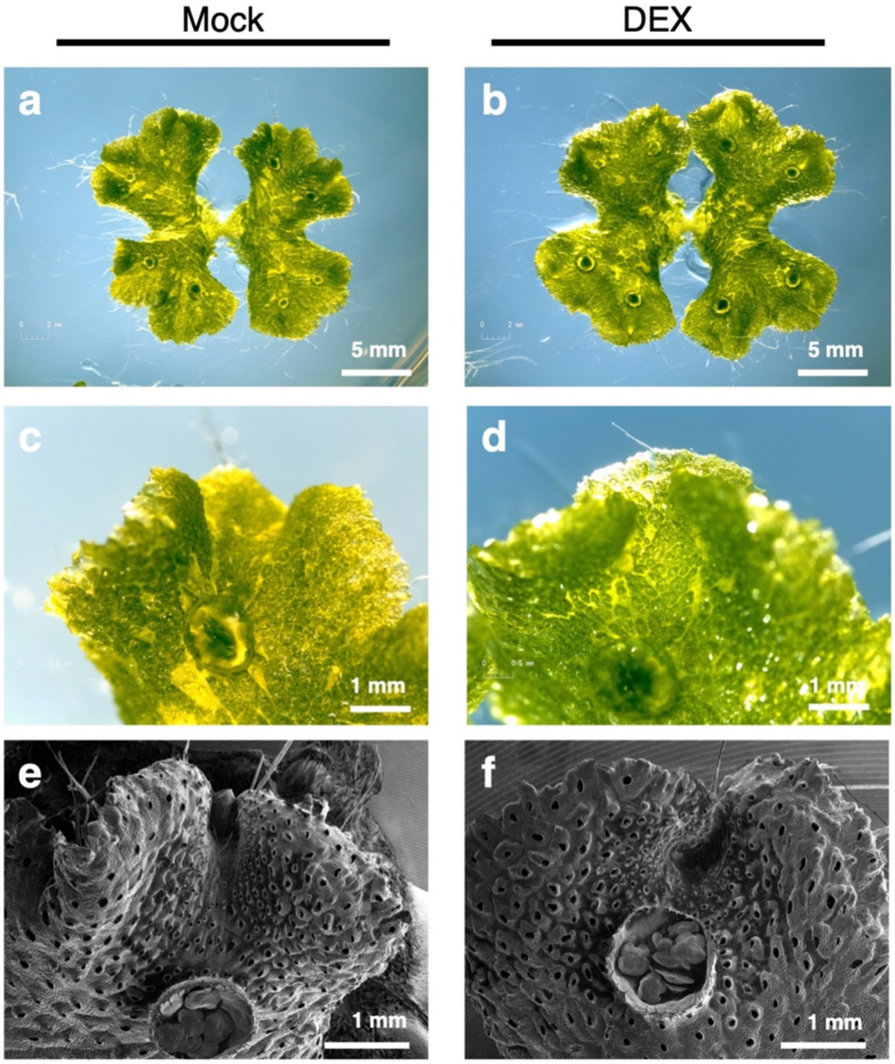


**Supplementary Figure 5. No morphological phenotype is conferred by DEX treatment in wild type thalli, related to Fig. 4.**
**(a and b)** 2-week-old thalli grown from tip of thalli of the wild type with **(b)** or without **(a)** 10 µM DEX treatment. **(c)** and **(d)** Close-up view of dorsal surface of (**a**) and (**b**), respectively. **(e)** and **(f)**, Scanning electron micrographs of wild type thalli with **(f)** or without (**e**) 10 µM DEX treatment**.**

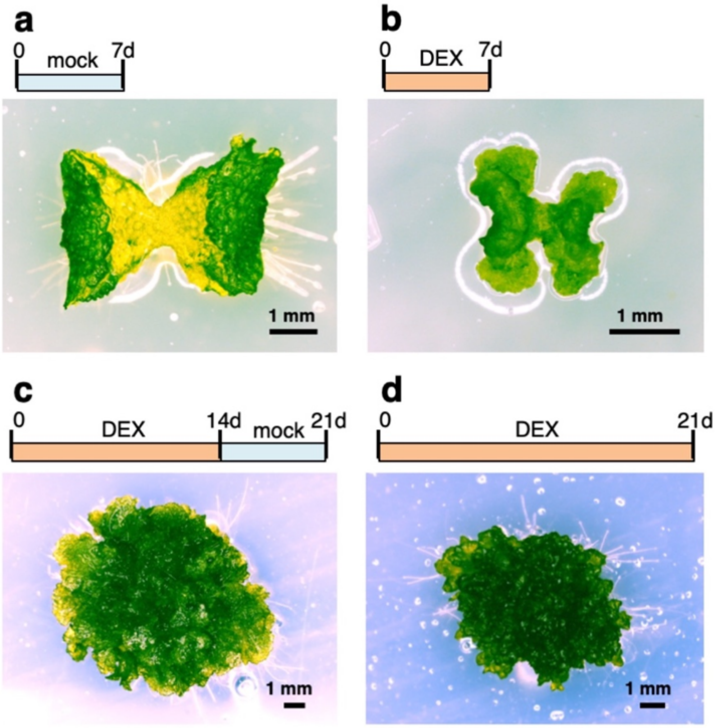


**Supplementary Figure 6. Induction of GCAM1 function suppresses growth and organ development in thallus, related to Fig. 4.**

**(a)** 1-week-old *MpEFpro:GCAM1-GR* transgenic plant treated with mock. **(b)** 2-week-old *MpEFpro:GCAM1-GR* transgenic plant treated with DEX. **(c)** 3-week-old *MpEFpro:GCAM1-GR* transgenic plant treated with DEX for the first 14 d and mock for the latter 7d. **(d)** 3-week-old *MpEFpro:GCAM1-GR* transgenic plant treated with DEX for 21d.

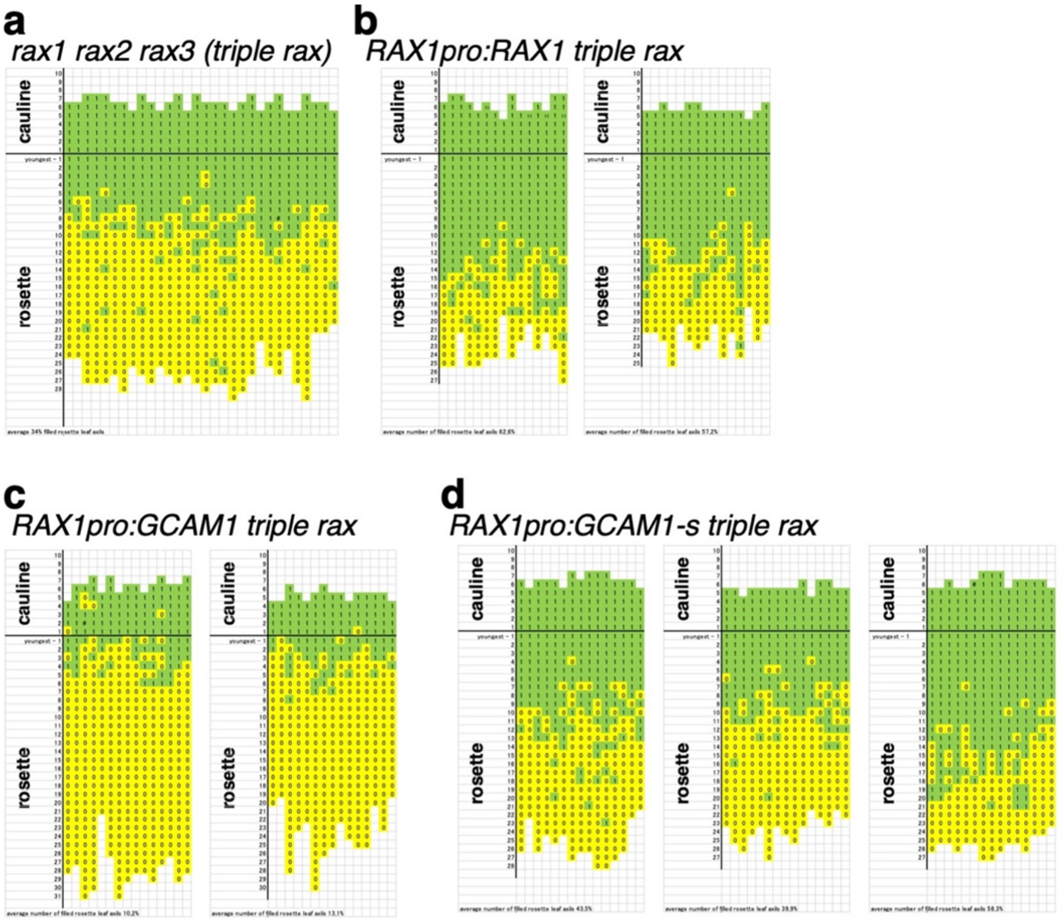


**Supplementary Figure 7. Interaction of GCAM1 with Arabidopsis mechanism for axillary meristem formation, related to Fig. 5.**

Schematic representation of axillary bud formation in leaf axils of Arabidopsis *rax* triple mutant (*n* = 30) **(a)** transformed with *RAX1pro:RAX1* (*n* = 15 for each lines) **(b)** or *RAX1pro:GCAM1* (*n* = 15 for each lines) **(c)** or *RAX1pro:GCAM1-s* construct (*n* = 15 for each lines) **(d)**. The long horizontal line represents the border between the youngest rosette leaf and the oldest cauline leaf. Each column represent a single plant, and each square within a column represents an individual leaf axil. The bottom row represents the oldest rosette leaf axils, with progressively younger leaves above. Green indicates the presence of an axillary bud, and yellow indicates the absence of an axillary bud in any particular leaf axil. Numbers in each square indicate number of axillary bud(s) in axil of each leaf. Arabidopsis plants were induced to flower under long-day conditions (16 h light at 22ºC and 8 h darkness at 18ºC) after 4 weeks under short-day conditions (8 h light at 22ºC and 16 h darkness at 18 ºC).

**Supplementary Table 1. Summary of the RNA-sequencing experiment**

| Sample name | Total reads counts | Aligned reads counts | | Alignment rate (%) |
|---|---|---|---|---|
| | | exactly once | >1 times | |
| Tak1; thallus 7d; rep1 | 18,318,924 | 16,558,099 | 621,394 | 90.4 |
| Tak1; thallus 7d; rep2 | 18,284,886 | 17,102,045 | 544,457 | 93.5 |
| Tak1; thallus 7d; rep3 | 19,162,536 | 18,006,202 | 569,092 | 94.0 |
| Tak1; gemma-cup 21d; rep1 | 17,216,306 | 15,496,802 | 604,487 | 90.0 |
| Tak1; gemma-cup 21d; rep2 | 14,489,889 | 13,625,766 | 421,670 | 94.0 |
| Tak1; gemma-cup 21d; rep3 | 15,883,627 | 14,812,606 | 473,619 | 93.3 |
| Tak1; mid-rib without gemma-cup 21d; rep1 | 14,774,027 | 13,622,665 | 427,230 | 92.2 |
| Tak1; mid-rib without gemma-cup 21d; rep2 | 16,075,894 | 15,127,377 | 452,720 | 94.1 |
| Tak1; mid-rib without gemma-cup 21d; rep3 | 15,312,864 | 14,337,977 | 474,028 | 93.6 |

**Supplementary Table 2. Transcription factor genes upregulated in gemma cup**

| Gene ID | TAIR10 BLAST tophit | | | | RPKM value (Mean) | | |
|---|---|---|---|---|---|---|---|
| | TF family | TOP hit in TAIR database | AGI code | E-value | Thallus | Mid-rib | Gemma cup |
| Mapoly0086s0035 | B3 | ABA INSENSITIVE 3 (ABI3) | AT3G24650 | 1E-34 | 8.4 | 6.7 | 25.3 |
| Mapoly0012s0202 | bHLH | RETARDED GROWTH OF EMBRYO 1 (RGE1) | AT1G49770 | 2E-14 | 4.0 | 1.5 | 14.6 |
| Mapoly0039s0003 | bHLH | RHD SIX-LIKE 1 (RSL1) | AT5G37800 | 4E-26 | 8.7 | 3.7 | 16.6 |
| Mapoly0073s0051 | bHLH | HECATE1 (HEC1) | AT5G67060 | 2E-20 | 10.1 | 6.4 | 44.2 |
| Mapoly0126s0029 | bHLH | RETARDED GROWTH OF EMBRYO 1 (RGE1) | AT1G49770 | 4E-28 | 1.9 | 1.0 | 4.0 |
| Mapoly0072s0050 | bZIP | ABSCISIC ACID RESPONSIVE ELEMENTS-BINDING FACTOR 2 (ABF2) | AT1G45249 | 2E-28 | 8.1 | 11.3 | 37.6 |
| Mapoly0061s0011 | C2H2 | ZINC FINGER PROTEIN 11 (ZFP11) | AT2G42410 | 3E-13 | 2.9 | 1.0 | 9.6 |
| Mapoly0008s0029 | R2R3-MYB | AtMYB117, LATERAL ORGAN FUSION 1 (LOF1) | AT1G26780 | 6E-56 | 6.2 | 5.6 | 16.5 |
| Mapoly0034s0034 | R2R3-MYB | REGULATOR OF AXILLARY MERISTEMS3 (RAX3) | AT3G49690 | 3E-59 | 0.8 | 6.3 | 82.1 |
| Mapoly0090s0091 | MYB-related | MYB DOMAIN PROTEIN 20 (MYB20) | AT1G66230 | 2E-13 | 26.4 | 15.5 | 111.5 |
| Mapoly0091s0012 | MYB-related | MYB FAMILY TRANSCRIPTION FACTOR | AT5G41020 | 4E-17 | 0.4 | 0.2 | 2.9 |

**Supplementary Table 3. Primers used in this study**.

| Name | Sequence (5'→3') | Usage |
|---|---|---|
| GCAM1-cds-I-F | GATGTCGGGACGCAAATAAT | qRT-RCR |
| GCAM1-cds-I-F | TGTCATTGCGTAGGGAGATTC | qRT-RCR |
| MpEF1a-F | TCACTCTGGGTGTGAAGCAG | qRT-RCR control |
| MpEF1a-R | GCCTCGAGTAAAGCTTCGTG | qRT-RCR control |
| GCAM1_ISH_probe_F | TGGTGGACGAGCATCAACAA | *in situ* RNA hybridization |
| GCAM1_ISH_probe_R* | tgcgtaatacgactcactatagggCGTGCGACTGGGAAGTAAGT | *in situ* RNA hybridization |
| MpHIS4_ISH_probe_F | GCCAAGCGTCATAGGAAGGT | *in situ* RNA hybridization |
| MpHIS4_ISH_probe_R* | tgcgtaatacgactcactatagggCCCGAACCCGTACAGAGTTC | *in situ* RNA hybridization |
| GCAM1-pro-L1 | caccCTCATGTACGCAATGGTTGG | Construction of GCAM1pro:GUS |
| GCAM1-pro-R2 | TCTGCAGCTCTTTCCGCAGCG | Construction of GCAM1pro:GUS |
| GCAM1-KI_5IF_L** | CCTAAGGTAGCGATTAATCGGACCCCCTGTACTGTAGAGTAC | Construction of GCAM1-Citrine knock-in construct |
| GCAM1-KI_5IF_R** | GGAGCCTCCAAGCTTAATACTGATCAATTTGGACAGGAAAGAATC | Construction of GCAM1-Citrine knock-in construct |
| GCAM1-KI_3IF_L** | TTATGTTTAAACTAGTGGGAGGAATGAAGGCCTGCTTGGTC | Construction of GCAM1-Citrine knock-in construct |
| GCAM1-KI_3IF_R** | TACCCTGTTATCCCTAGGTACCTTCAGACGGTTTCGCC | Construction of GCAM1-Citrine knock-in construct |
| d-F | GTACAACCGCTCTCAGGACCTATCT | For checking of gene targeting site for GCAM1-Citrine |
| d-R | GAACCCTAAGCTTTGGAGCTTGATA | For checking of gene targeting site for GCAM1-Citrine |
| e-F | GCCGGAACCTGAATAGAAAGAAAGAAG | For checking of gene targeting site for GCAM1-Citrine |
| e-R | AGATGAACTTCAGGGTCAGCTTGC | For checking of gene targeting site for GCAM1-Citrine |
| H1F | GTATAATGTATGCTATACGAAGTTATGTTT | For checking of gene targeting site for GCAM1-Citrine |
| f-R | ATGACAAGAAGAAGAAATGTCACAGTCG | For checking of gene targeting site for GCAM1-Citrine |
| GCAM1_5IF_L** | CTAAGGTAGCGATTAAAAGATAGGTCGCTTCGGTCAG | Construction of the targeting vector for GCAM1 knock-out |
| GCAM1_5IF_R** | GCCCGGGCAAGCTTAGATGGCATCCACATTCGATAAA | Construction of the targeting vector for GCAM1 knock-out |
| GCAM1_3IF_L** | TAAACTAGTGGCGCGTGATCACTGGGAACAAACGAAG | Construction of the targeting vector for GCAM1 knock-out |
| GCAM1_3IF_R** | TTATCCCTAGGCGCGTCGGTTAAAGCCTATCACACCA | Construction of the targeting vector for GCAM1 knock-out |
| a-F | CCTGTAATGAGTGGATTCGCTTG | For checking of gene targeting site for $gcam1^{KO}$ |
| c-R | TAGGACTCGATGCTGAACTCGTC | For checking of gene targeting site for $gcam1^{KO}$ |
| b-F | GCAAACCGTGAAACTCCTGTAAG | For checking of gene targeting site for $gcam1^{KO}$ |
| b-R | CTTTGGAGTCTTTCCGGGTTATG | For checking of gene targeting site for $gcam1^{KO}$ |
| P1R | GAAGGCTTCTGATTGAAGTTTCCTTTTCTG | For checking of gene targeting site for $gcam1^{KO}$ |
| H1F | GTATAATGTATGCTATACGAAGTTATGTTT | For checking of gene targeting site for $gcam1^{KO}$ |
| GCAM1_tg1-L*** | gcacccagcctctcgGTCCTTGGTCGCCCGAAGgtttagagctagaa | Construction of the genome editing vector for GCAM1: target 1 |
| GCAM1_tg1-R*** | ttctagctctaaaacCTTCGGGCGACCAAGGACcgagaggctgggtgc | Construction of the genome editing vector for GCAM1: target 1 |
| GCAM1_tg2-L*** | gcacccagcctctcgTACAGGTCTGAAGCGCTGgtttagagctagaa | Construction of the genome editing vector for GCAM1: target 2 |
| GCAM1_tg2-R*** | ttctagctctaaaacCAGCGCTTCAGACCTGTAcgagaggctgggtgc | Construction of the genome editing vector for GCAM1: target 2 |
| CRISPR-check_L | CCTGTAATGAGTGGATTCGCTTG | For checking of the edited site for GCAM1 |
| CRISPR-check_R | TAGGACTCGATGCTGAACTCGTC | For checking of the edited site for GCAM1 |
| GCAM1-cds-L1 | caccATGCAACCTTTGCCCACTAA | Construction of entry clone containing GCAM1 coding region |
| GCAM1-cds-nsR | ACTGATCAATTTGGACAGGAAAGA | Construction of entry clone containing GCAM1 coding region |
| GCAM1-cds-sR | TTAACTGATCAATTTGGACAGGA | Construction of entry clone containing GCAM1 coding region |
| Myb37-pro-SbfIF | CATCCTGCAGGATAAATTAAACTATCGATTGGGTC | Introduction of GCAM1 into A. thaliana rax triple mutant |
| Myb37-pro-AscIR | CATGGCGCGCCTTCTCTCGTTAGTGAATTGAAGT | Introduction of GCAM1 into A. thaliana rax triple mutant |
| GFPAscIF | GGCGCGCCATGAGTAAAGGAGAAGAACT | Introduction of GCAM1 into A. thaliana rax triple mutant |
| GFPpolylinkerR | GAGCTCCCTAGGTTATTTGTATAGTTCATCCA | Introduction of GCAM1 into A. thaliana rax triple mutant |
| RAX1AscIF | CATGGCGCGCCATGGGAAGAGCACCGTGTTG | Introduction of GCAM1 into A. thaliana rax triple mutant |
| RAX1SacIR | CATGAGCTCCTAGGAGTAGAAATAGGGCA | Introduction of GCAM1 into A. thaliana rax triple mutant |
| GCAM1AscIF | CATGGCGCGCCATGCAACCTTTGCCCACTAA | Introduction of GCAM1 into A. thaliana rax triple mutant |
| GCAM1AvrIIR | CGTCCTAGGTTAACTGATCAATTTGGACA | Introduction of GCAM1 into A. thaliana rax triple mutant |
| GCAM1NewAscIF | CATGGCGCGCCATGGGTCGAGCTCCATGTTG | Introduction of GCAM1 into A. thaliana rax triple mutant |
| GCAM1AvrIIR | CGTCCTAGGTTAACTGATCAATTTGGACA | Introduction of GCAM1 into A. thaliana rax triple mutant |

*Bases in lower case indicate a priming site of T7 RNA polymerase

**Underlined bases indicate the homologous sequences for cloning into pJHY-TMp1-Cit or pJHT-TMp1 using Gibson Assembly (NEB) or In-fusion cloning (TAKARA) kit.

***Bases in lower case indicate the homologous sequences for cloning into pMpGE_En01 using In-gusion cloning (TAKARA) kit.
